# Architectural Mediator subunits are differentially essential for global transcription in yeast

**DOI:** 10.1101/2020.03.02.973529

**Authors:** Jason P. Tourigny, Kenny Schumacher, Didier Devys, Gabriel E. Zentner

## Abstract

The modular Mediator complex is a coactivator of RNA polymerase II transcription. We show that depletion of the main complex scaffold Med14 or the head module scaffold Med17 is lethal and results in global transcriptional downregulation in yeast, though Med17 removal has a markedly greater negative effect. Depletion of Med14 or Med17 impairs pre-initiation complex (PIC) assembly similarly, suggesting that the differential transcriptional effects observed are not due to differing extents of defective PIC formation. Co-depletion of Med14 and Med17 reduced transcription and TFIIB promoter occupancy similarly to Med17 ablation alone, suggesting that the independent head module can weakly stimulate transcription *in vivo*, though not to a level that maintains viability. We suggest that, while the structural integrity of complete Mediator and the head module are both important for PIC assembly, the head module additionally promotes optimal PIC function and is thus the key functional module of Mediator in this regard.

## Introduction

Mediator is a conserved, essential regulatory complex that transmits regulatory information between transcription factors bound to distal regulatory elements and promoter-bound general transcription factors (GTFs) and RNA polymerase II (RNAPII). The 25 subunits of budding yeast Mediator are classified into four modules: head, middle, tail, and kinase. Med14 acts as a scaffold, holding together the head, middle, and tail modules via extensive protein-protein interactions (Plaschka et al., 2015; Robinson et al., 2015; Tsai et al., 2014). The head module contacts RNAPII and its formation is nucleated by Med17 (Imasaki et al., 2011). The head and middle modules form the functional core Mediator (cMed), an assembly of 15 mostly essential subunits (Liu et al., 2001; Plaschka et al., 2015). The nonessential tail and kinase modules serve a regulatory role, influencing the recruitment and function of cMed (Jeronimo et al., 2016).

The majority of data on the role of Mediator in transcription has been obtained using *srb4-138*, a temperature-sensitive (ts) allele of Med17 that dissociates the head module from the remainder of the complex at elevated temperature (Linder et al., 2006). Single-gene and global mRNA measurements from this strain following Med17 inactivation revealed a decrease comparable to that seen for a ts RNAPII mutant (Holstege et al., 1998; Thompson and Young, 1995), though such measurements cannot distinguish the effects of altered mRNA synthesis and decay. More recent studies have demonstrated global decreases in nascent RNA synthesis (Plaschka et al., 2015) and RNAPII occupancy (Paul et al., 2015) in *srb4-138* cells at elevated temperature, but analysis of Mediator function under heat shock is complicated by the general decrease in RNAPII occupancy observed at elevated temperature (Warfield et al., 2017) and the involvement of Mediator in heat shock transcription (Anandhakumar et al., 2016; Kim and Gross, 2013; Kremer et al., 2012). Furthermore, dominant mutations in the *MED6* and *MED22* genes, both encoding head subunits, suppress *srb4-138* lethality at the restrictive temperature but do not compensate for *SRB4* deletion (Lee et al., 1998), suggesting that the mutant protein retains some function at elevated temperature.

Nuclear depletion of Mediator subunits with anchor-away (AA) (Haruki et al., 2008) has also been used to probe their roles in transcription. As Med17 nucleates the head module (Imasaki et al., 2011), it stands to reason that its depletion would reduce or ablate head module formation. Med17 AA is not lethal and results in a relatively modest global reduction in RNAPII occupancy (Bruzzone et al., 2018; Petrenko et al., 2017). Similarly, Med14 AA leads to a moderate reduction in RNAPII occupancy at ∼400 well transcribed genes and, surprisingly, maintains viability (Petrenko et al., 2017). It has thus been proposed that Mediator modules contribute independently to overall complex function and can individually promote substantial transcription such that viability is maintained. Given that deletion of Med14 or Med17 is lethal (Sakai et al., 1990; Thompson et al., 1993) while AA of these subunits is not, it is likely that AA yields inducible hypomorphic alleles that are useful for revealing sets of genes heavily dependent on Mediator but may not allow for determination of the full effects of individual Mediator subunit loss. Indeed, auxin-inducible degron (AID)-mediated depletion of Med14 decreases RNAPII occupancy of nearly all genes (Warfield et al., 2017).

Given these disparate results, we sought to gain a better understanding of the relationship between architectural integrity of Mediator and transcription *in vivo*. We directly measured transcription by labeling, purification, and sequencing of newly synthesized RNA (nsRNA) following AID-mediated ablation of the main complex scaffold Med14 or the head module nucleator Med17. Depletion of either factor was lethal and globally downregulated transcription. Removal of Med17 caused a substantially greater downregulation of transcription than depletion of Med14 despite comparable reductions in PIC component binding to promoters, arguing that differential effects on PIC formation and/or stability are not the underlying cause of the distinct transcriptional phenotypes observed. Simultaneous depletion of Med14 and Med17 resulted in transcriptional downregulation equivalent to that seen with removal of Med17 alone, arguing that the head module can modestly stimulate transcription independent of complete Mediator. Our results indicate that cohesion of complete Mediator and the head module are both essential for transcription at a level that supports viability in yeast, but the degrees to which they are necessary for transcription are different. We propose that both aspects of Mediator integrity are equally important for PIC assembly but that the head module is additionally required for optimal PIC function.

## Results and Discussion

### Functional impairment of Mediator with the auxin degron system

We used the AID system (Nishimura et al., 2009) to inducibly deplete the structurally essential subunits Med14 and Med17. Based on available biochemical and structural data, we reasoned that Med14 loss would dissociate Mediator into disconnected but stable head, middle, and tail modules (Plaschka et al., 2015; Robinson et al., 2015; Tsai et al., 2014); we thus use the term ‘Mediator dissociation’ in reference to Med14 depletion. To ablate the head module, we targeted Med17, which acts as a scaffold for head module assembly (Imasaki et al., 2011). Depletion of Med14-AID and Med17-AID was very efficient, with near-complete removal observed by 30 min after addition of the auxin indole-3-acetic acid (3-IAA) (Figure 1A). We also generated a strain in which Sua7/TFIIB was tagged with AID, reasoning that this would allow us to compare the transcriptional effects of Mediator subunit depletion with those of directly impaired PIC assembly and function. Sua7-AID displayed similar depletion kinetics to those seen for Med14-AID and Med17-AID (Figure 1A). The Med14-AID, Med17-AID, and Sua7-AID strains all failed to grow on YPD medium containing 3-IAA, while the parental *Os*TIR1-expressing strain exhibited normal growth (Figure 1B). Given that Med14 and Med17 have been shown to be essential by classical genetic approaches (Sakai et al., 1990; Thompson et al., 1993), these results argue that the AID system is suitable for full functional genomic analysis of essential Mediator subunits.

**Figure 1.**
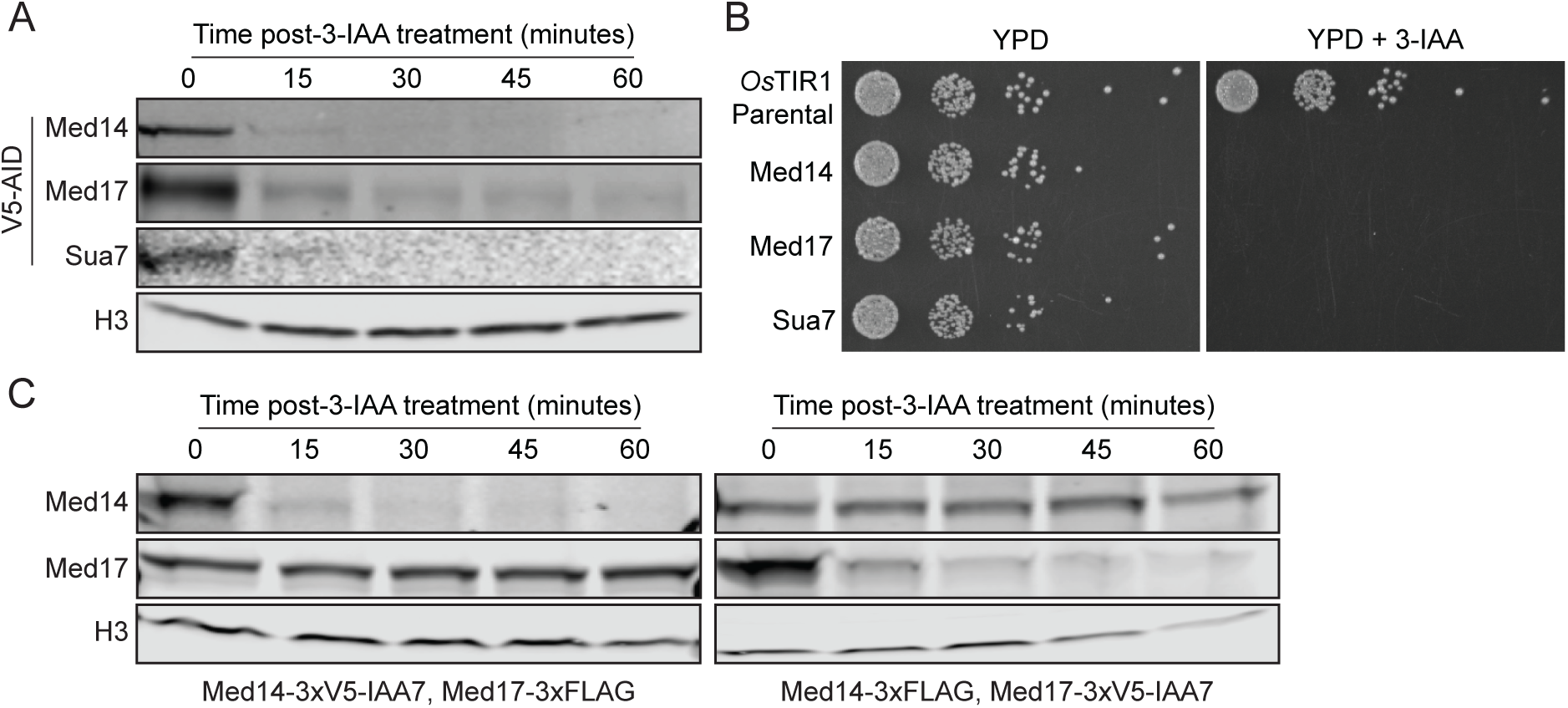
Destabilization of complete Mediator and the Mediator head module with the auxin degron system. (A) Western blots showing the kinetics of 3-IAA-mediated depletion of AID-tagged factors. (B) Spot assays assessing growth of AID-tagged strains on YPD plates containing DMSO or 500 µM 3-IAA. (C) Western blots showing levels of Med17 following Med14 depletion and Med14 levels following Med17 depletion.

We next sought to validate the specificity of our approach for individual Mediator subunits. To this end, we tagged Med17 with a 3xFLAG epitope in the Med14-AID background and vice versa. We observed no effect of Med14 depletion on Med17 stability even after a 1 h 3-IAA treatment (Figure 1C), suggesting that AID targeting of Med14 does not nonspecifically affect Med17. Similarly, removal of Med17 had no impact on Med14 protein levels (Figure 1C). Thus, AID appears to target specific Mediator modules, in contrast to previous studies using AA where Med14 AA removes Med17 from the nucleus (Anandhakumar et al., 2016) and vice versa (Petrenko et al., 2017). Notably, we observed with both the AID- and FLAG-tagged forms of each factor that Med17 levels were substantially higher than those of Med14. This variation is not due to differences in sample processing and imaging, as the sets of Med14-AID/Med17-FLAG and Med14-FLAG/Med17-AID samples were electrophoresed on a single gel and blotted and exposed on a single membrane to ensure comparability (Figure S1). While we cannot rule out differences in western blot transfer efficiency as the cause of the apparent differences in Med14 and Med17 abundance we observe, these results are consistent with values recently reported in an integrative analysis of nearly two dozen quantitative protein abundance datasets (Ho et al., 2018), with mean values of 2,138 Med14 and 2,873 Med17 molecules per cell and median values of 1,989 Med14 molecules and 3,342 Med17 molecules/cell.

### Mediator dissociation and head module ablation downregulate global transcription to different extents

To assess the acute transcriptional effects of various forms of Mediator impairment, we performed 4-thiouracil (4tU) labeling, biotinylation, purification, and high-throughput sequencing (4tU-seq) of nsRNA. To establish a transcription-null baseline, we first assessed total and nsRNA levels following Sua7 depletion. As we observed strong depletion of all factors by 30 min of 3-IAA treatment (Figure 1A), we chose this early time point for 4tU labeling to avoid the potentially confounding effects of increasing cell dysfunction and death. *S. cerevisiae* cells were labeled for 6 min following 3-IAA treatment and spiked with a defined fraction of similarly labeled *S. pombe* cells to enable normalization against an exogenous reference. After sequencing, we quantified the levels of 5,080 genes encoding verified ORFs and RNAPII-dependent sn/snoRNAs in steady-state (total) and nsRNA fractions. The defect in transcription caused by Sua7 depletion was evident even in the total RNA fraction: at a fold change (FC) cutoff of 2 and an adjusted p-value threshold of 0.05, 4,129 genes were significantly downregulated, with a median FC of -4.87 for all genes (Figure 2A). Analysis of nsRNA levels following Sua7 removal revealed a massive downregulation of transcription, with 5,007 genes significantly downregulated and a median FC of -16.61 for all genes (Figure 2B).

**Figure 2.**
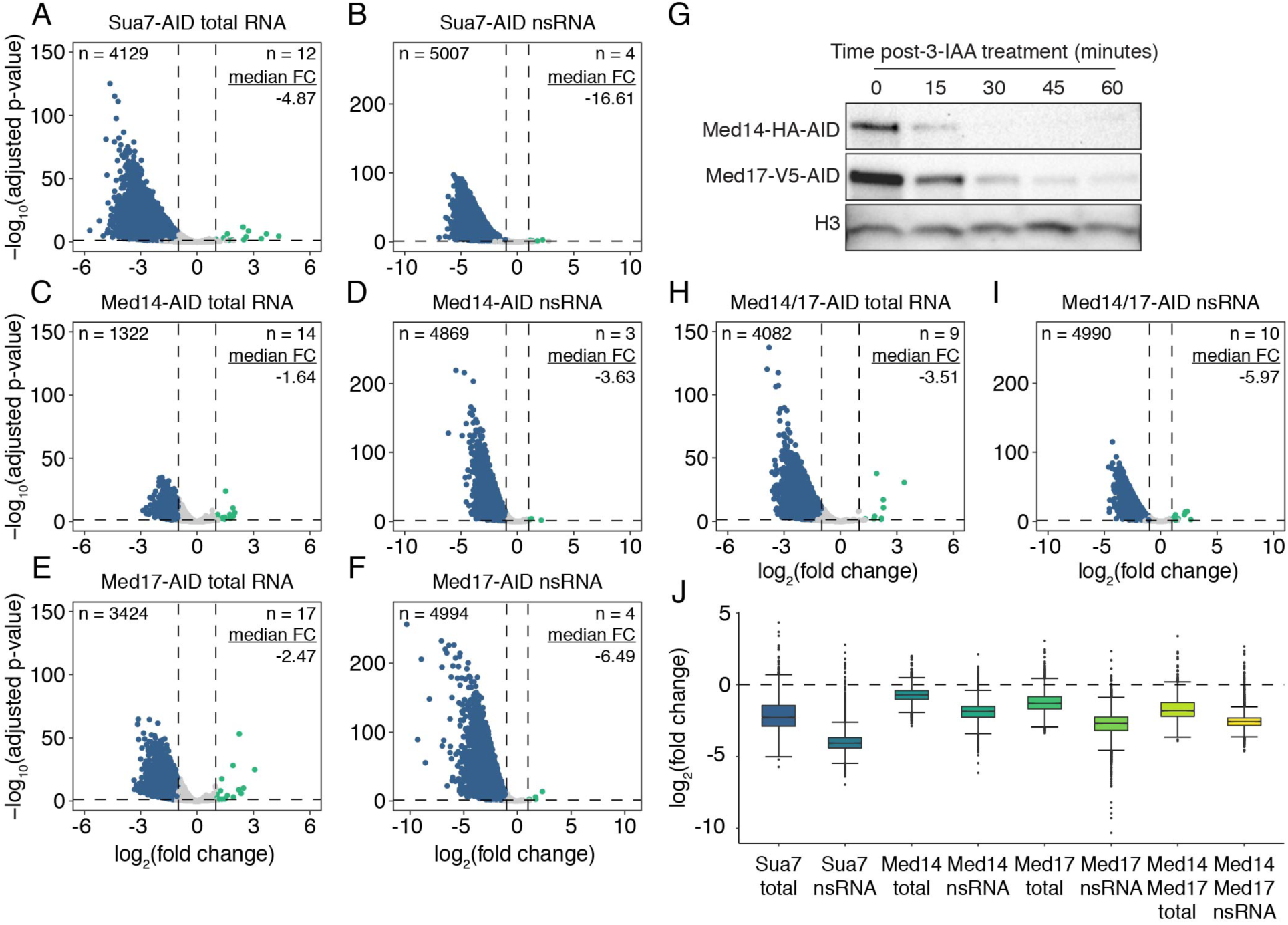
Architectural Mediator subunit depletion results in global transcriptional downregulation. (A-B) Volcano plots comparing the fold changes and adjusted p-values of 5,080 RNAPII-transcribed genes in the (A) total and (B) nsRNA fractions after Sua7 depletion. A fold change cutoff of 2 with an adjusted p-value threshold of 0.05 was used to assess significance. Significantly downregulated genes are colored blue and significantly upregulated genes are colored teal. Numbers of significantly downregulated and upregulated genes as well as the median linear fold change of all genes are given. (C-D) Same as (A-B) but for Med14 depletion. (E-F) Same as (A-B) but for Med17 depletion. (G) Western blots showing the kinetics of 3-IAA-mediator depletion of Med14 and Med17 in the dual degron strain. (H-I) Same as (A-B) but for Med14/17 co-depletion. (J) Boxplots summarizing the fold changes of all 5,080 analyzed genes in the total and nsRNA fractions from each experiment. Whiskers represent the smallest and largest values up to IQR * 1.5.

Having established a baseline of transcription ablation via Sua7 depletion, we next determined the transcriptional consequences of Med14 and Med17 removal. Med14 loss resulted in a modest reduction of steady-state transcript levels, with 1,322 genes displaying reduced abundance and a median FC of -1.64 for all genes (Figure 2C). In contrast, analysis of nsRNA after Med14 depletion revealed a robust global downregulation of transcription, with 4,869 genes significantly downregulated and a median FC of -3.63 for all genes (Figure 2D). Notably, Med17 ablation resulted in a much stronger effect on steady-state transcript levels than that of Med14, with 3,424 genes significantly downregulated and a median FC of -2.47 for all genes (Figure 2E). The impact of Med17 loss on nsRNA levels was also substantially more pronounced, with 4,994 genes significantly downregulated at a median FC of -6.49 for all genes (Figure 2F). From these data, we conclude that the Mediator complex as a whole and the head module are both essential for global transcription at levels supporting viability, though the effect of ablating the head module is more dramatic. However, depletion of neither Med14 nor Med17 reduced nsRNA levels to the extent seen following Sua7 ablation. These observations support a model in which transcription occurs from PICs lacking Mediator *in vivo*. However, because Med14 or Med17 depletion is lethal, the transcription that does persist is insufficient to support life (Figure 1B).

### Simultaneous depletion of Med14 and Med17 is equivalent to loss of Med17 with respect to global transcription

Our results thus far indicate that depletion of Med17 has greater negative effect on transcription than that of Med14 loss. We therefore speculated that the head module, which is a biochemically stable entity (Imasaki et al., 2011; Takagi et al., 2006), might remain associated with the PIC following dissociation from the middle and tail modules and impart some residual level of transcriptional stimulation. Were this the case, we would expect that simultaneous depletion of Med14 and Med17 would result in transcriptional downregulation comparable to that observed with the destruction of Med17 alone. We therefore generated a strain bearing both Med14-AID and Med17-AID (Figure 2G) and sequenced total and nsRNA following 3-IAA treatment. We observed a greater decrease in total RNA levels with Med14/17 co-depletion than with either single Mediator degron (Figure 2H). However, at the level of nsRNA, Med14/17 depletion had a slightly milder negative effect than single Med17 depletion (Med14/17-AID median FC = -5.97, Med17-AID median FC = -6.49) with a similar number of genes downregulated (Med14/17-AID, 4,990 genes; Med17-AID, 4,994 genes) (Figure 2I). For all factors tested, we observed a substantially greater decrease in nsRNA than total RNA levels (Figure 2J), indicating buffering of steady-state transcript levels via decreased mRNA decay (Sun et al., 2012).

While our data suggest that the continued association of the head module with the PIC following Med14 depletion results in a modest degree of transcriptional stimulation, the potential mechanism is unclear. Recombinant head module only weakly stimulates transcription *in vitro* (Takagi et al., 2006) and does not restore transcriptional activity to Mediator-depleted nuclear extract (Plaschka et al., 2015). Furthermore, structural rearrangements of Mediator in the context of the RNAPII holoenzyme appear to rely on the coherence of the complete complex (Tsai et al., 2014). Alternatively, the tail module may promote transcription in the absence of Med14, as it has been proposed that a triad of tail subunits (Med2, Med3, and Med15) may promote transcription when severed from cMed by removal of the tail/middle connector subunit Med16 (Galdieri et al., 2012; Zhang et al., 2004). However, microarray analysis of transcript levels in *med16*Δ yeast suggests limited transcriptional changes comparable to those observed with mutations in members of the tail triad or kinase module (Ansari et al., 2012; Kemmeren et al., 2014; van de Peppel et al., 2005). It is therefore unlikely that the independent tail could provide global compensation of transcriptional downregulation following Med14 depletion. In sum, while our data support the idea that Mediator modules can play roles outside the context of the complete complex *in vivo*, they indicate that these functions are unable to support transcription sufficient for viability in the absence of intact Mediator or the head module (Figure 1B).

### Gene-specific effects of Mediator impairment

We next sought to determine the transcriptional effects of middle- and head-specific Mediator destabilization on specific gene categories. Recent work analyzing the effects of acute depletion of SAGA and TFIID subunits led to a proposed reclassification of yeast genes as coactivator-redundant (CR) or TFIID-dependent (Donczew et al., 2020). CR genes are modestly sensitive to acute depletion of subunits of either SAGA or TFIID but are dramatically affected by simultaneous depletion of subunits from both complexes, while TFIID-dependent genes are heavily dependent on TFIID but show little change in transcription following rapid depletion of SAGA subunits. Med17 depletion resulted in greater downregulation of both classes of genes than did depletion of Med14, though the degrees of downregulation were more comparable for CR than for TFIID-dependent genes (Figure 3A). Consistent with our analysis of all genes (Fig. 2J), co-depletion of Med14 and Med17 resulted in slightly milder downregulation of CR and TFIID-dependent genes than did ablation of Med17 alone (Figure 3A). We observed a similar effect for genes classified as SAGA or TFIID-dominated (Huisinga and Pugh, 2004) (Figure 3B).

**Figure 3.**
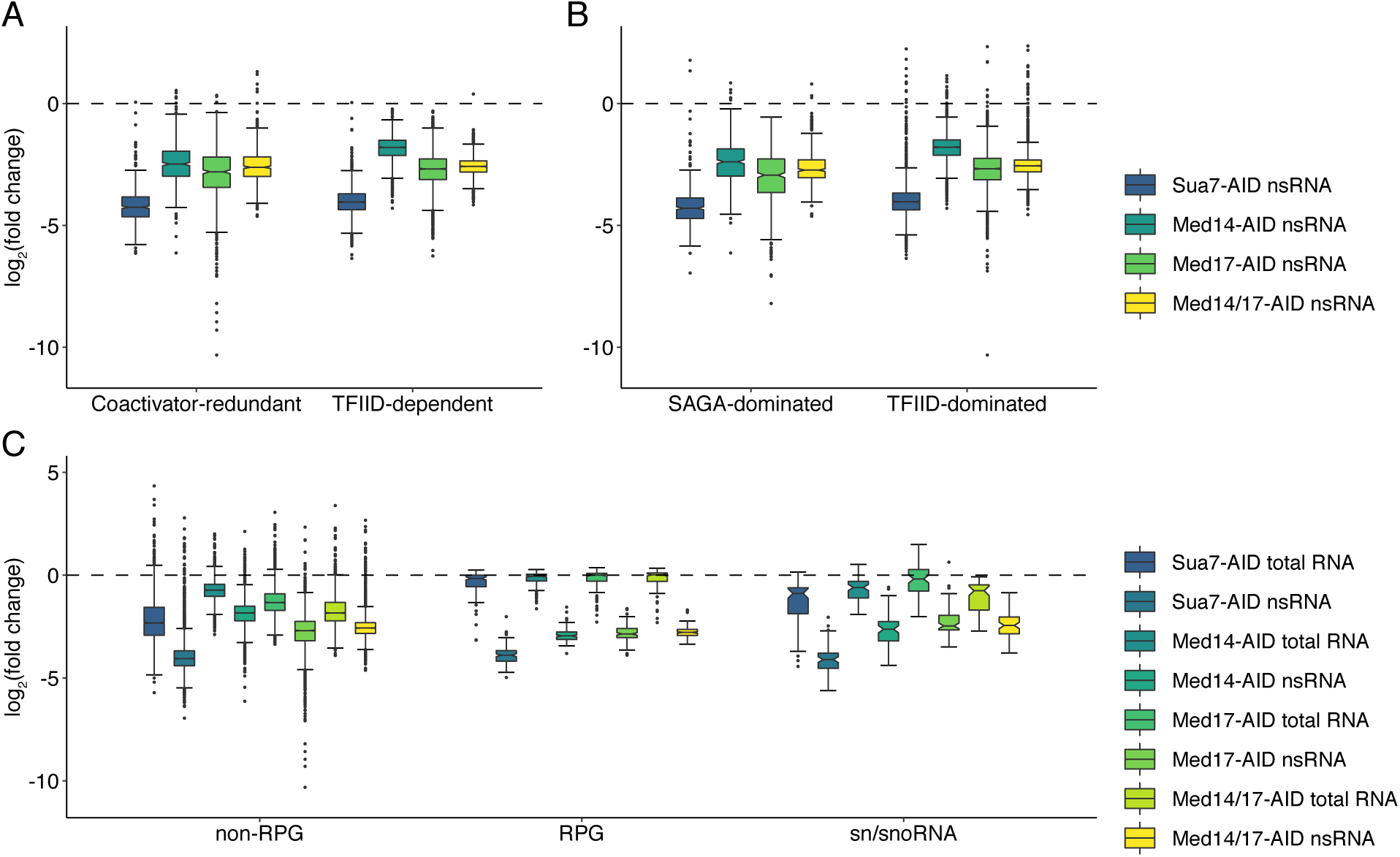
Gene-specific effects of Mediator subunit depletion. (A-B) Boxplots of log_2_(fold changes) in nsRNA levels following Sua7, Med14, Med17, or Med14/17 depletion for genes classified as (A) coactivator redundant or TFIID-dependent or (B) SAGA or TFIID-dominated (exclusive of RPGs). (C) Boxplots of log_2_(fold changes) in total and nsRNA levels following Sua7, Med14, Med17, or Med14/17 depletion for all non-RPG protein coding genes, RPGs, and sn/snoRNAs. Whiskers represent the smallest and largest values up to IQR * 1.5. Notches represent 1.58 * IQR / sqrt(n), which equates to a ∼95% confidence interval for comparing medians.

Our observation that Med14 removal preferentially downregulates CR and SAGA-dominated genes is consistent with previous data using RNAPII ChIP-seq to interrogate transcription following Med14 AA (Petrenko et al., 2017). The distinct effects of Med14 removal on CR or SAGA-dominated genes and TFIID-dependent genes may be due to differences in regulatory region structure. The promoters of genes more reliant on SAGA for full expression tend to contain consensus TATA boxes (Basehoar et al., 2004), and the UASs of TATA-containing promoters tend to be located further upstream than at TATA-less promoters (Erb and van Nimwegen, 2011). As increased distance between a UAS and a TATA box in the context of a reporter gene attenuates its expression (Dobi and Winston, 2007) and deletion of Mediator tail subunits affects distance-dependent UAS activity (Reavey et al., 2015), it stands to reason that the connection between the head and tail modules may be more important at SAGA-reliant genes based on their greater UAS-TATA distance. However, our data contrast with previous studies on the preference of Med17 for certain categories of genes. Two independent studies using Med17 AA reported a preferential impact on RNAPII occupancy of SAGA dominated genes (Bruzzone et al., 2018; Petrenko et al., 2017), while our data show that Med17 is required equally regardless of coactivator dependence.

### Med14 and Med17 are equally important for ribosomal protein gene and sn/snoRNA transcription

We next analyzed the effects of Mediator impairment on the transcription of ribosomal protein genes (RPGs), a co-regulated set of 137 genes encoding the structural components of the ribosome (Warner, 1999), and small nuclear and nucleolar RNAs (sn/snoRNAs), which we recently found to be dependent on Med14 for full expression (Tourigny et al., 2018). In contrast to what we observed for genes classified by coactivator dependence, Med14 and Med17 depletion reduced RPG and sn/snoRNA transcription to comparable extents (Figure 3C). Strikingly, we also observed that, while RPG transcription was heavily downregulated by Sua7 or Mediator subunit depletion, steady-state RPG transcript levels were essentially unchanged, in stark contrast to non-RPG protein-coding transcripts (Figure 3C). We observed a similar phenomenon for sn/snoRNAs, though to a lesser extent. These observations suggest that RPG transcripts and potentially sn/snoRNAs are highly buffered against decreased synthesis. In the case of RPGs, this buffer is likely due to their high codon optimality (Presnyak et al., 2015), but it is unclear if this phenomenon might be more general. We therefore examined the relationship of optimal codon usage with mean nsRNA levels across all DMSO-treated samples. This analysis revealed a highly significant positive correlation (Spearman’s *ρ* = 0.53, p < 2.2 × 10^−16^) (Figure S2), consistent with previous comparisons of RNA synthesis rates and codon optimality in budding and fission yeast (Harigaya and Parker, 2016) as well as functional studies of the relationship between transcription and codon optimality in *Neurospora crassa* (Zhou et al., 2016).

### Complete and head-specific Mediator ablation affect PIC formation to comparable extents

Given that a primary function of Mediator is promotion of PIC assembly (Soutourina, 2018), we next assessed the effects of Med14 and Med17 depletion on promoter occupancy of the PIC by ChIP-seq. We constructed derivatives of the Med14-AID and Med17-AID strains with a 3xFLAG-tagged allele of a representative subunit of several GTFs: Sua7 (TFIIB), Tfa2 (TFIIE), and Tfg2 (TFIIF). We then visualized enrichment of each GTF around all gene promoters, regardless of expression level, as heatmaps. Depletion of either Med14 or Med17 resulted in equivalent decreases in Sua7 occupancy at promoters, with more highly bound promoters displaying greater loss of binding (Figure 4A-B, S3). We observed the same pattern with Tfa2 and Tfg2 (Figure 4C-F). We also assessed Sua7 binding to the genome following co-depletion of Med14 and Med17. The magnitude of the reduction in Sua7 binding following simultaneous Med14 and Med17 removal was comparable to that seen with single depletion of either Med14 or Med17 (Figure 4G). Thus, depletion of Med14 or Med17 results in comparable decreases in PIC association with promoters but drastically different degrees of transcriptional downregulation.

**Figure 4.**
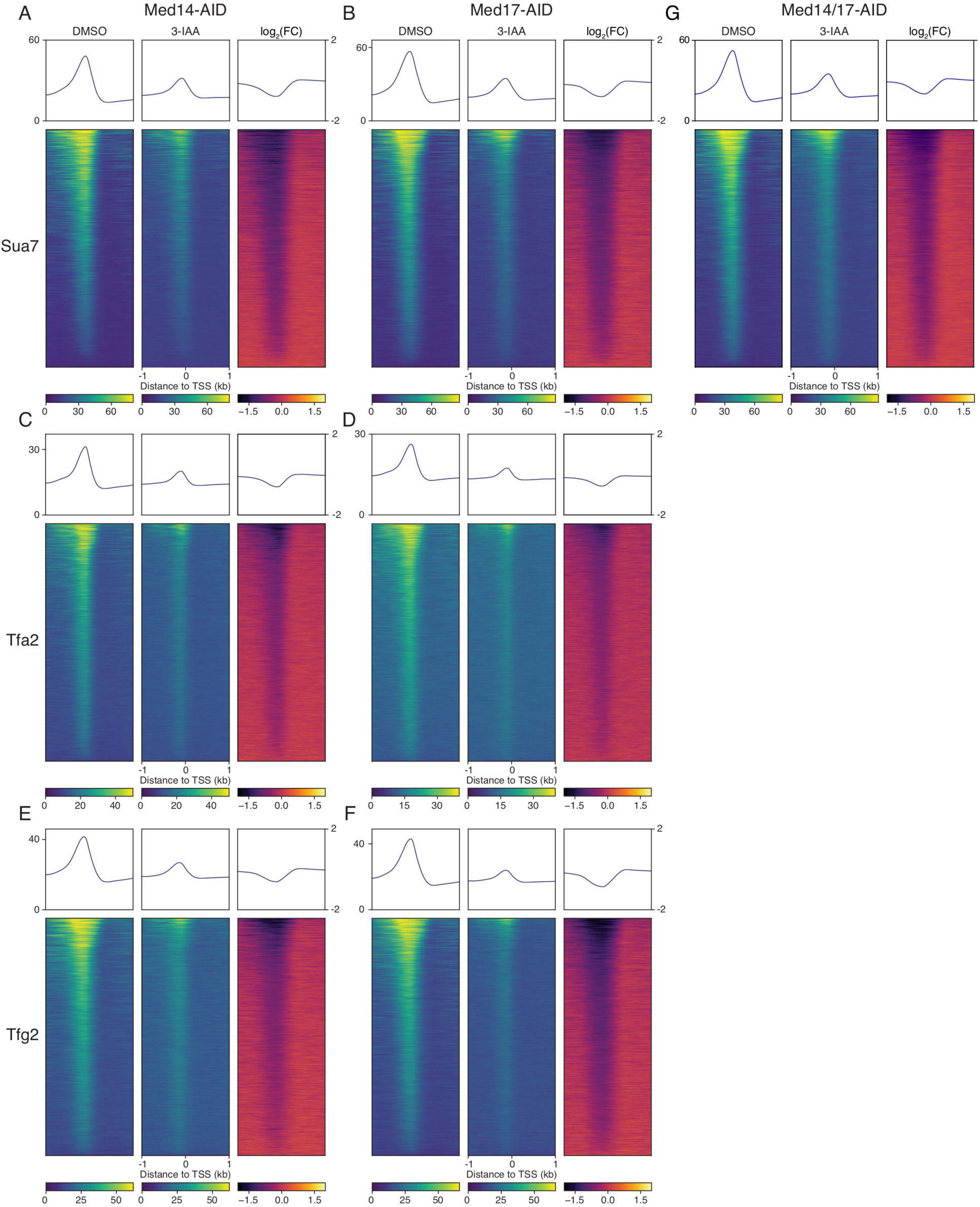
Equivalent effects of Med14, Med17, and Med14/17 depletion on GTF promoter binding. (A-F) Average plots and heatmaps of (A-B) Sua7, (C-D) Tfa2, and (E-F) Tfg2 ChIP-seq signal around TSSs following DMSO or 3-IAA treatment of the indicated strain. (G) Same as in (A-F) but for Sua7 ChIP-seq in the Med14/17-AID strain. The three heatmaps in each panel are sorted descending by average signal in the DMSO-treated sample.

From these observations, it appears that, while intact Mediator is necessary for their complete function and assembly at promoters, PICs persist at some level following acute depletion of essential Mediator components, at least within the 30 min timeframe assessed here. This observation is consistent with previous work employing AA of head module subunits (Jeronimo and Robert, 2014) and our present finding that depletion of Sua7, a *bona fide* GTF, resulted in a substantially higher degree of global transcriptional downregulation than loss of either Med14 or Med17. However, the decreases in GTF binding we observed were comparable for Med14 and Med17 depletion, indicating that the substantial difference in the degree of transcriptional downregulation resulting from depletion of these factors is not simply due to differential effects on PIC formation and/or stability.

### Concluding remarks

Our results provide clear evidence that coherence of complete Mediator and the head module are both essential for viability-supporting transcription, but indicate substantial differences in the contributions of these facets of Mediator structural integrity to global transcription in yeast. Depletion of Med17, expected to dissociate the head module, has a significantly greater negative impact on transcription than removal of Med14, required for association of the head, middle, and tail modules. We also found that, while Med14 ablation preferentially affects transcription of CR genes, Med17 removal results in equivalent downregulation of genes regardless of their annotated coactivator dependence. The transcriptional defects observed with both Med14 and Med17 depletion are also less severe than those seen following direct impairment of PIC formation via depletion of Sua7. This suggests that PICs lacking functional Mediator are still transcriptionally active; however, as yeast depleted of Med14 or Med17 fail to grow, this residual transcription is insufficient for viability. Notably, depletion of either Med14 or Med17 caused a comparable decrease in promoter association of several GTFs, arguing that differential effects on PIC assembly do not completely explain the markedly different transcriptional phenotypes we observed. Lastly, our data suggest that the head module can promote transcription independent of the larger Mediator complex and thus support the notion that individual Mediator modules can play complex-independent roles; however, they also indicate that these roles are likely to be minor and insufficient to maintain viability. Based on the substantially greater decrease in transcription caused by Med17 versus Med14 removal, the lack of gene class specificity in the negative transcriptional effects of Med17 depletion, and the comparable effects of depleting either factor on PIC association with promoters, we propose that the head module acts as the major functional module of Mediator *in vivo* by promoting optimal PIC function to stimulate transcription.

## Materials and methods

### Yeast methods

Med14, Med17, and Sua7 were tagged with 3xV5-IAA7 using pL260 (pFA6a-3xV5-IAA7-kanMX6) (Miller et al., 2016) in the GZY191 (BY4705 pGPD-OsTIR1-HIS3) background (Tourigny et al., 2018). Med14 was tagged with 3xHA-IAA7 using pGZ364 (pFA6a-3xHA-IAA7-URA3). Mediator and PIC components were tagged with 3xFLAG using pFA6a-6xGLY-3xFLAG-hphMX4 (Funakoshi and Hochstrasser, 2009) (a gift from Mark Hochstrasser, Addgene plasmid #20755) or a derivative thereof in which the hphMX4 marker is replaced with a *TRP1* cassette (pGZ392). For spot assays, strains were grown overnight and diluted to an OD_600_ of 1.0. Tenfold serial dilutions were plated on YPD with and without 500 µM 3-IAA and plates were imaged after 48 h of growth at 30°C. Strain genotypes are provided in Table S1.

### 4tU-seq

RNA labeling and purification was performed as previously described (Baptista and Devys, 2018; Baptista et al., 2017). Sequencing libraries were constructed by the GenomEast platform at the Institut de Génétique et de Biologie Moléculaire et Cellulaire using the Illumina TruSeq Stranded Total RNA LT Sample Prep Kit. Libraries were sequenced on the on the Illumina HiSeq 4000 platform with the following parameters: 50 cycles paired-end (Med14-AID and Med17-AID), 100 cycles paired-end (Sua7-AID), 50 cycles single-end (Med14/17-AID).

### ChIP-seq

ChIP was performed as described (Tourigny et al., 2018). Sequencing libraries were constructed by the Indiana University Center for Genomics and Bioinformatics (CGB) using the NEBNext Ultra II Library Prep Kit for Illumina and sequenced for 38 or 80 cycles in paired-end mode on the Illumina NextSeq 500 platform. Sua7 ChIP-seq data in the Med14-AID background with DMSO or 3-IAA treatment was previously published (Tourigny et al., 2018) (GSE112721).

### Data analysis

#### Total RNA-seq/4tU-seq

Reads were mapped to a chimeric genome composed of the *S. cerevisiae* (sacCer3) and *S. pombe* (ASM294v2) genomes using STAR (Dobin et al., 2013) v2.5.3a. Quantification was performed with uniquely aligned reads using htseq-count (Anders et al., 2014) v0.6.1p1 with annotations from Ensembl version 94 for *S. cerevisiae* and Ensembl fungi version 41 for *S. pombe* and union mode. *S. cerevisiae* read counts were normalized across samples with the size factors computed by the median-of-ratios method (Anders and Huber, 2010) on *S. pombe* genes to make these counts comparable between samples. Differential expression analysis was performed with DESeq2 (Love et al., 2014) v1.16.1. Log_2_(fold changes), p-values, and adjusted p-values for the 5,080 analyzed transcripts are given in Table S2. Codon usage of the 5,011 mRNAs analyzed here was performed with coRdon (Elek et al., 2019) and the fraction of optimal codons for each transcript was calculated using a previously published list of optimal codons (Presnyak et al., 2015). Optimal codon percentage was then correlated with the average of spike-in-normalized nsRNA counts divided by median transcript length in kb across all DMSO-treated samples. Data analysis and visualization were performed with the R programming language. Code used for figure generation is available upon request.

#### ChIP-seq

Paired-end ChIP-seq data were aligned to the sacCer3 genome build using Bowtie2 (Langmead and Salzberg, 2012) with default settings plus ’-I 10 -X 700 --no-unal --dovetail --no-discordant - -no-mixed’. SAM files were converted to sorted BAMs using SAMtools (Li et al., 2009). CPM-normalized bigWig files were generated using deepTools (Ramírez et al., 2016) v3.3.0 bamCoverage with 1 bp bins and counts per million (CPM) normalization. Log_2_(3-IAA/DMSO) bigWig files were generated using deepTools bamCompare with 1 bp bins and signal extraction scaling (SES) normalization (Diaz et al., 2012). Matrices of ChIP-seq signal around TSSs to be plotted as heatmaps were generated with deepTools computeMatrix using reference-point mode, 1 bp bins, and 1000 bp windows upstream and downstream of each TSS. Heatmaps were then generated with deepTools plotHeatmap.

## Data availability

Total RNA-seq/4tU-seq and ChIP-seq data have been deposited in GEO (accession pending).

## Supporting information

Figures S1-S3, Table S1, and legend for Table S2

## Author contributions

JPT, KS, DD, and GEZ designed the study. JPT performed all experiments except 4tU-seq, which was performed by KS. JPT, KS, and GEZ analyzed data. JPT and GEZ wrote the manuscript with input from all authors.

## Acknowledgements

We thank Sebastian Grünberg, Steven Hahn, Laszlo Tora, and members of the Zentner Lab for helpful discussions throughout the course of this work. This work was supported by Agence Nationale de la Recherche grant ANR-18-CE12-0026 SAGA-Retina to DD and National Institutes of Health grant R35GM128631 to GEZ.

