## Supplementary material for "Architectural Mediator subunits are differentially essential for global transcription in yeast": Figures S1-S3, Table S1, and legend for Table S2

Contents:

Figures S1-S3
Table S1
Legend for Table S2

Figure 1A
Single Degrons

Med14-V5-IAA7 (top), H3 (bottom)
merged from LI-COR

Med17-V5-IAA7 (top), H3 (bottom)

Sua7-V5-IAA (top), H3 (bottom)

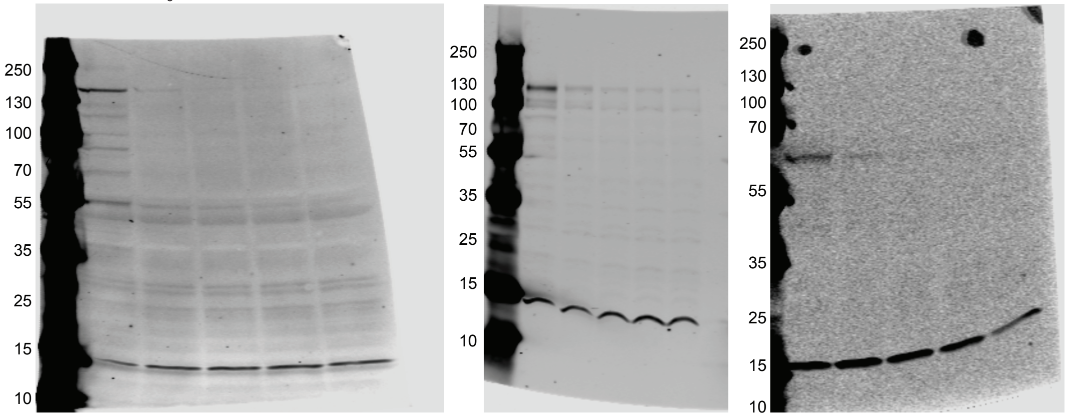


Figure 1C
Single Degrons
w/ inverse FLAG

MED14-V5-IAA7 (top left), MED17-V5-IAA7 (top right), H3 (bottom)
These are from the same LI-COR scan, with contrast turned up for MED14

MED14-FL (top left), MED17-FL (top right), H3 reprobe (far bottom)
Same double degron timecourse samples as to the left, but probed for FLAG first
Contrast turned down for MED17 and up for MED14 similarly

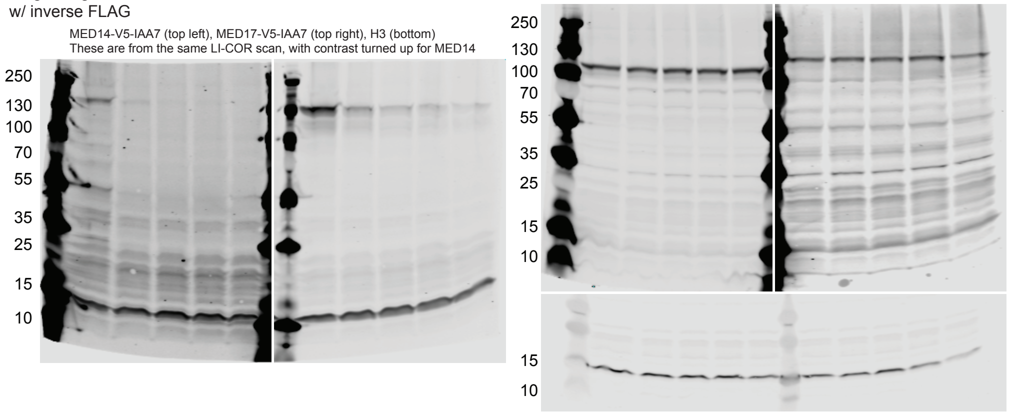


Figure 2G
Double Degron

300s chemiluminescent exposure for Med14-HA-IAA7, top right

35s for Med17-V5-IAA7, top left

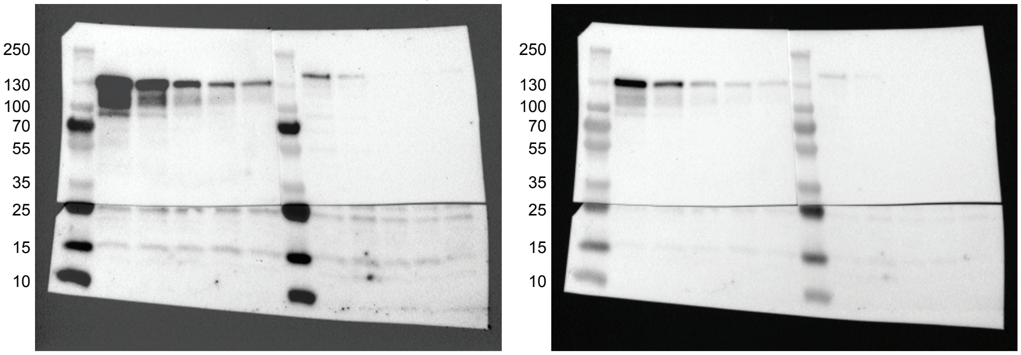


6s for H3 reprobe

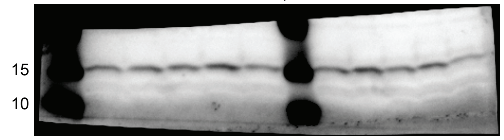


**Figure S1. Full western blot scans**

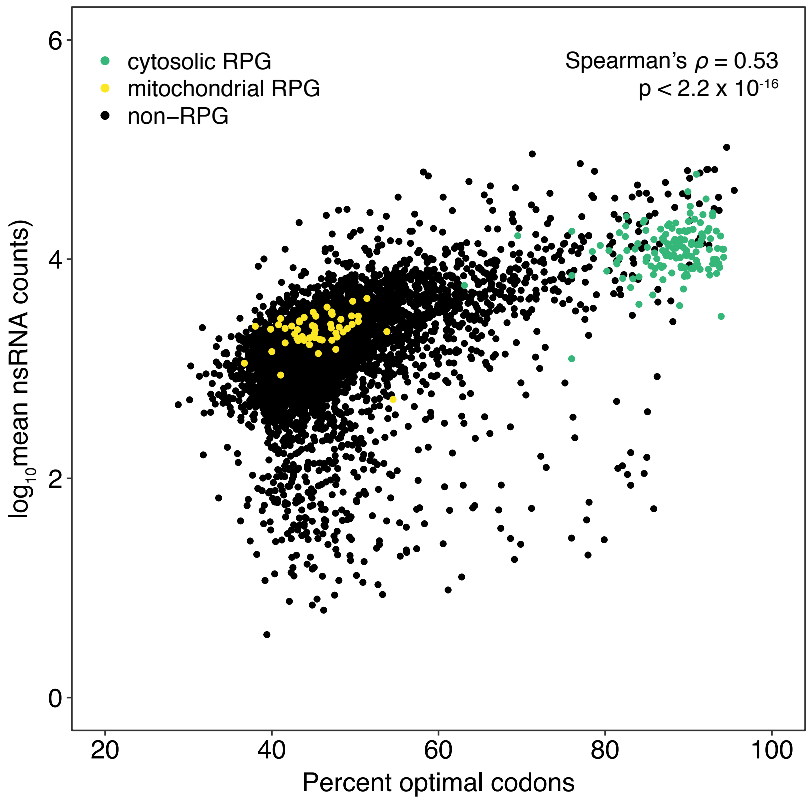


**Figure S2. Comparison of codon optimality and transcription**

Scatterplot comparing the optimal codon percentage and average normalized nsRNA levels of mRNA genes (n = 5,011) across all DMSO-treated samples. Optimal codons were obtained from a previous study (Presnyak et al., 2015). Cytosolic and mitochondrial RPGs are highlighted as control gene groups with high and low codon optimality, respectively.

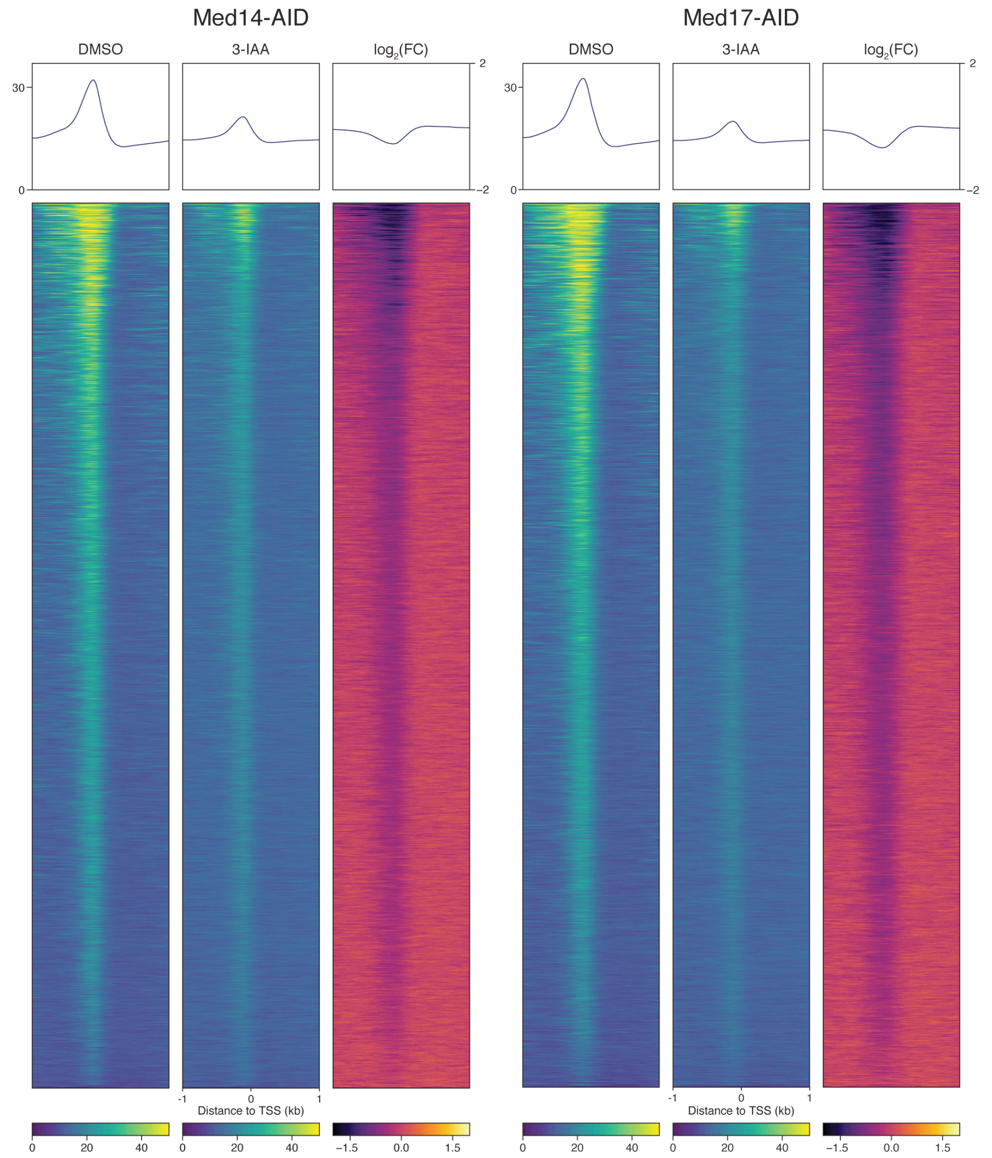


**Figure S3. Sua7 ChIP-seq replicate analysis**

Average plots and heatmaps of Sua7 ChIP-seq signal around TSSs following DMSO or 3-IAA treatment of the (A) Med14-AID or (B) Med17-AID strain. An average plot and heatmap of the $log_2$(3-IAA/DMSO) ChIP-seq signal is also shown in each panel.

| Strain | Genotype | Source |
|---|---|---|
| DHP43 (*S. pombe*) | *h-* | D. Devys |
| GZY191 | *MATα ade2Δ::hisG his3Δ200 leu2Δ0 lys2Δ0 met15Δ0 trp1Δ63 ura3Δ0 his3::pGPD1-OsTIR1-HIS3* | (Tourigny et al., 2018) |
| GZY219 | GZY191 *MED14-3V5-IAA7-kanMX6* | (Tourigny et al., 2018) |
| GZY269 | GZY191 *MED17-3V5-IAA7-kanMX6* | This work |
| GZY272 | GZY269 *SUA7-6GLY-3FLAG-hphMX4* | This work |
| GZY279 | GZY219 *TFA2-6GLY-3FLAG-hphMX4* | This work |
| GZY282 | GZY269 *TFA2-6GLY-3FLAG-hphMX4* | This work |
| GZY337 | GZY219 *MED14-6GLY-3FLAG-TRP1* | This work |
| GZY339 | GZY219 *TFG2-6GLY-3FLAG-TRP1* | This work |
| GZY341 | GZY269 *MED17-6GLY-3FLAG-TRP1* | This work |
| GZY361 | GZY269 *TFG2-6GLY-3FLAG-TRP1* | This work |
| GZY364 | GZY191 *SUA7-3V5-IAA7-kanMX6* | This work |
| GZY380 | GZY269 *MED14-3HA-IAA7-URA3* | This work |
| GZY388 | GZY380 *SUA7-6GLY-3FLAG-TRP1* | This work |

**Table S1. Yeast strains used in this study**

All *S. cerevisiae* strains were constructed in the BY4705 background.

**Table S2. Information on total RNA-seq and 4tU-seq results**

This Excel spreadsheet contains mean $\log_2$(fold changes), p-values, and adjusted p-values for the 5,080 genes analyzed in total RNA-seq and 4tU-seq experiments. Gene classification information (e.g. coactivator dependence) is also provided.
